# Ancient mitochondrial DNA pathogenic variants putatively associated with mitochondrial disease

**DOI:** 10.1101/2020.05.13.094243

**Authors:** Draga Toncheva, Dimitar Serbezov, Sena Karachanak-Yankova, Desislava Nesheva

**Author notes:** Corresponding author: Prof. Draga Toncheva Department of Medical Genetics, Faculty of Medicine Medical University of Sofia, Bulgaria Dean Office, Zdrave str. 2, 1431 Sofia, Bulgaria.

## Abstract

Mitochondrial DNA variants associated with diseases are widely studied in contemporary populations, but their prevalence has not yet been investigated in ancient populations. The publicly available AmtDB database contains 1443 ancient mtDNA Eurasian genomes from different periods. The objective of this study was to use this data to establish the presence of pathogenic mtDNA variants putatively associated with mitochondrial diseases in ancient populations. The clinical significance, pathogenicity prediction and contemporary frequency of mtDNA variants were determined using online platforms. The analyzed ancient mtDNAs contain six variants designated as being “*confirmed pathogenic*” in modern patients. The oldest of these, m.7510T>C in the *MT-TS1* gene, was found in a sample from the Neolithic period dated 5800-5400 BCE. All six have well established clinical association, and their pathogenic effect is corroborated by very low population frequencies in contemporary populations. In addition, ten variants designated as possibly or likely pathogenic were detected. The oldest of these were two variants in the *MT-TD* gene, m.7543A>G and m.7554G>A, from Neolithic samples dated 8205-7700 BCE. A novel mutation in contemporary populations, m.4440G>A in the *MT-TM* gene, is established in 12 ancient mtDNA samples from different periods ranging from 2800 BCE to 920 CE. The pathogenic effect of these possibly/likely pathogenic mutations is not yet well established, and further research is warranted. All detected mutations putatively associated with mitochondrial disease in ancient mtDNA samples are in tRNA coding genes. Most of these mutations are in a mt-tRNA type (Model 2) that is characterized by loss of D-loop/T-loop interaction. Seven mutations are located in CS-Anticodon stem, 4 are located in AS-Acceptor stem, 2 in TS-TΨC stem, and single mutations are found in DL-Dihydrouridine Loop, CL-Anticodon Loop and DS-Dihydrouridine stem. Exposing pathogenic variants in ancient human populations expands our understanding of their origin.

## Introduction

The scarcity of prehistoric human remains hampers obtaining complete picture of disorder incidence in ancient times. Altered or affected bones in skeletal remains might provide information about certain diseases, such as cancers (1, 2) and rheumatic diseases (3). A lesion on an archaic *Homo* mandible from Kanam, Kenya (Middle to Late Pleistocene) (4) and a fibrous dysplasia on a Neanderthal rib (older than 120000 years) from the site of Krapina, Croatia (5) are early confirmation of neoplastic disease. Neoplastic tumors have however been detected in early *Homo* samples as old as 1.7 million years ago, and these provide further insight into the outset of human cancers (6). Mummified human remains of a 5300-year-old Neolithic man (Ötzi, The Tyrolean Iceman) show hardening of the arteries, suggesting predisposition for coronary heart disease (7). Based on information on subsistence, geography and sample age, Berens *et al* (2017) estimate the genetic disease risk for 3180 loci in 147 ancient genomes, and find it to be similar to that of modern day humans (8). Focusing on individual genomes, however, they estimate that the overall genomic health of the Altai Neanderthal is worse than 97% of present day humans and that Ötzi the Tyrolean Iceman had a genetic predisposition to gastrointestinal and cardiovascular diseases (8).

Data on the prehistoric origin of mitochondrial diseases is however notably lacking. The first disease linked directly to mtDNA mutation was discovered in 1988 (9). Recently, whole-genome sequencing of mtDNA has led to significant advances in our understanding of mitochondrial diseases. Rare pathogenic mutations in mitochondrial DNA cause monogenic mitochondrial diseases involving multiple systems and are associated with variable clinical phenotypes. The severity of the clinical and biochemical phenotype caused by pathogenic mtDNA mutations has however been found to be roughly proportionate to the percent mutant heteroplasmy (10, 11). Nucleotide polymorphisms accumulate in mtDNA during human evolution forming mitochondrial haplogroups, and these also might alter the penetrance of mitochondrial diseases (12). Specific subclades of haplogroup J, for example, have been shown to affect the penetrance and pathogenicity of Leber’s hereditary optic neuropathy (13). Certain mtDNA mutations and haplogroups are also predictors of both lifespan and risk of various age-associated disease, including degenerative diseases, cancer, diabetes, heart failure, sarcopenia and Parkinson’s disease (14).

We had previously performed whole-genome sequencing on 25 Thracian mtDNA samples dated 3000-2000 BCE, and 608 mtDNA variants were detected (15). Only one of these, however, m.15326A>G (rs2853508) is designated as likely pathogenic, associated with familial breast cancer (16). This variant was found in all analyzed by us samples, and MitoMap (2019, update nr.3) database estimates 0.98 population frequency (17). Such high frequency leads to the conclusion that this variant is common and probably not disease related.

The objective of this study was to investigate pathogenic mutations in ancient mtDNA and thus to provide further insight into the emergence of mitochondrial diseases.

## Materials & Methods

We used the comprehensive data of the ancient mtDNA genome sequences from the Ancient mtDNA database (18) which compiles human mitochondrial variation studies in ancient populations. We analyzed the *fasta* files of 1443 samples from different periods: Paleolithic 10, Mesolithic 96; Neolithic 341; Copper 242; Bronze 368; Iron 152 and Middle Ages 234, analyzed by whole genome sequencing. The total number of variants detected in these samples was 3191.

We used various publicly available databases to gather information to identify mtDNA sequence variants and to gather information about each variant, including its clinical significance and contemporary population frequencies:

- MitoMap (2019, update nr.3) is a human mitochondrial genome database that contains 14383 SNVs, 49135 full-length sequences and 72235 control region sequences (17).

MitoMap classifies a variant as being “confirmed pathogenic” if it meets the set of criteria outlined by Mitchell et al 2006 (19), Yarham et al 2011 (20), Wong 2007 (21) and Gonzalez-Viogue et al 2014 (22): (1) independent reports of two or more unrelated families with evidence of similar disease; (2) evolutionary conservation of the nucleotide (for RNA variants) or amino acid (for coding variants); (3) presence of heteroplasmy; (4) correlation of variant with phenotype / segregation of the mutation with the disease within a family; (5) biochemical defects in the OXPHOS genes constituent complexes I, III, or IV in affected or in multiple tissues; (6) functional studies showing differential defects segregating with the mutation (cybrid or single fiber studies); histochemical evidence of a mitochondrial disorder; and (8) for fatal or severe phenotypes, the absence or extremely rare occurrence of the variant in large mtDNA sequence databases.

- MITOMASTER was used to identify, annotate and evaluate the potential biological significance of nucleotide variants (23).
- GenBank database provides access to the most up-to-date and exhaustive DNA sequence information, and was used to get information on variant frequencies in contemporary populations (24).
- MitoTIP is an *in silico* tool for predicting pathogenicity of novel mitochondrial tRNA variants (25). It integrates multiple sources of information, including the position of the variant within the tRNA, conservation across species and population frequencies, to provide a prediction for the likelihood that novel single nucleotide variants would cause disease.
- HmtVar uses algorithms to determine the importance of the variant position in tRNAs and was utilized to predict the pathogenicity and potential impact of mtDNA variants (26).
- Complementing the information obtained using the abovementioned tools, literature survey was performed on variants designated as “*confirmed pathogenic*|” in an effort to acquire comprehensive picture of the evidence for their disease causing effect.

## Results

Out of 3191 unique mtDNA variants established in the 1443 analyzed ancient samples, six are designated as being “*confirmed pathogenic*” and 10 as likely/possibly pathogenic by MitoMap. For each of these variants, we review the available evidence in HmtVar and in the scientific literature for their pathogenic effects.

### Confirmed pathogenic mutations

#### m.5703G>A (rs199476130)

The variant m.5703G>A was found in four ancient mtDNA samples, one from the Neolithic period, three from the Iron Age and one from the Middle Ages. The two Iron Age samples are from the same site in Russia, extracted from the remains of a male and a female, and these two individuals might have been related (27). This is also the pathogenic variant established in ancient mtDNA from archeological sites spanning the widest geographical range, i.e. from Poland to Mongolia (Figure 1). This mutation is in the *MT-TN* gene, which encodes tRNA Asparagine (Table 1). MitoMap and HmtVar designate this mutation as pathogenic.

**Table 1.** Mitochondrial DNA mutations established in ancient samples putatively associated with mitochondrial disease.

| Pathogenic mutations |  |  |  |  |  |  |  |
| --- | --- | --- | --- | --- | --- | --- | --- |
| Nr | Variants (17, 18)/ Gene product/ Strand (26) | Period/Years (18) | Country / Location / Sample | GB Freq. | Mito Tip classification | HmtVar Prediction | Mitochondrial Disease (MitoMap) |
| 1 | m.5703G>A (rs199476130) tRNA Asparagine /L | Neolithic (2880-2776 BCE) | Poland / Koszyce / RISE1170 (55) | 0.00% | Cfrm Path. | Pathogenic (M2-P27) | CPEO / MM / COX deficiency (Heteroplasmy) |
|  |  | Iron Age (900-600 BCE) | Russia / Grishkin Log 1 / DA4 (27) |  |  |  |  |
|  |  | Iron Age (156-134 BCE) | Mongolia / Omnogobi / DA45 (27) |  |  |  |  |
|  |  | Middle Ages (10-375 CE) | Poland / Kowalewko, Greater Poland / PCA0002 (56) |  |  |  |  |
| 2 | m.3243A>G (rs199474657) tRNA Leucine/H | Bronze Age (2029-1911 BCE) | Germany / Haunstetten, Postillionstr . / POST_16 (57) | 0.02% | Cfrm Path. | Pathogenic (M0-P14) | MELAS / LS / DMDF / MIDD / SNHL / CPEO / MM / FSGS / ASD / Cardiac+multi-organ dysfunction (Heteroplasmy) |
| 3 | m.5650G>A tRNA Alanine/L | Iron Age (900-600 BCE) | Russia / Grishkin Log 1 / DA5 (27) | 0.00% | Cfrm Path. | Pathogenic (M2-P6) | Myopathy (Heteroplasmy) |
|  |  | Iron Age (766-729 BCE) | Kazakhstan / Kurgan Borli, Osakarovskij / DA11 (27) |  |  |  |  |
|  |  | Iron Age (156-134 BCE) | Mongolia / Omnogobi / DA45 (27) |  |  |  |  |
| 4 | m.8340G>A, tRNA Lysine/H | Middle Ages (10-375 CE) | Poland / Kowalewko, Greater Poland / PCA0018 (56) | 0.00% | Cfrm Path | Pathogenic (M2-P51) | Myopathy/Exercise Intolerance/Eye disease+SNHL |
| 5 | m.14674T>C,<br>tRNA<br>Glutamic<br>acid/L | Bronze Age<br>(1592-1591<br>BCE) | Mongolia / Mitjurino / DA231<br>(27) | 0.01% | Cfrm Path | Pathogenic<br>(M2-P73) | Reversible COX deficiency<br>myopathy<br>(Homoplasmy) |
| 6 | m.7510T>C<br>(rs199474820)<br>tRNA Serine/L | Neolithic<br>(5800-5400<br>BCE) | Bulgaria / Malak Preslavets /<br>I1109 (58) | 0.00% | Cfrm Path | Pathogenic<br>(M1-P5) | SNHL<br>(Heteroplasmy) |
| **Likely/Possibly pathogenic** |  |  |  |  |  |  |  |
| 7 | m.1624C>T<br>tRNA<br>Valine/H | Middle Ages<br>(10-375 CE) | Poland / Kowalewko, Greater<br>Poland / PCA0037 (55) | 0.00% | PP | Pathogenic<br>(M2-P25) | Leigh Syndrome<br>(Homoplasmy) |
| 8 | m.4440G>A<br>tRNA<br>Methionine/H | Neolithic<br>(2800-2776<br>BCE) | Poland / Koszyce/ PCA0037 (55) | 0.00% | PP | Pathogenic<br>(M2-P42) | MM<br>(Heteroplasmy) |
|  |  | Iron Age (900-600 BCE) | Russia / Grishkin Log 1 / DA9<br>(27) |  |  |  |  |
|  |  | Iron Age (800-773 BCE) | Kazakhstan / Birlik, Kurgan 12,<br>Bajanaul DA17 (27) |  |  |  |  |
|  |  | Iron Age (600-400 BCE) | Kazakhstan / Karaganda, Kurgan<br>Sjartas, Shetskiy / DA10 (27) |  |  |  |  |
|  |  | Iron Age (357-342 BCE) | Kazakhstan / Kurgan Sjiderti 17,<br>Burial 1, Sjiderte, Pavlodar /<br>DA20 (27) |  |  |  |  |
|  |  | Iron Age (201-148 BCE) | Mongolia / Hovsgol, Grave #18 /<br>DA38 (27) |  |  |  |  |
|  |  | Iron Age (49<br>BCE-53 CE) | Mongolia / Arkhangai, Grave #1 /<br>DA39 (27) |  |  |  |  |
|  |  | Iron Age (47 BCE-24 CE) | Kazakhstan / Naurzum, Kurgan (3), Naurzumskijzapovednik DA30 (27) |  |  |  |  |
|  |  | Iron Age (177-190 CE) | Kazakhstan / Kurgan nr. 50, Japyryk / DA81 (27) |  |  |  |  |
|  |  | Iron Age (397-570 CE) | Kazakhstan / Kurgan nr. 1, Baskya 2 / DA385 (27) |  |  |  |  |
|  |  | Middle Ages (550-850 CE) | Kazakhstan / Bt, 2015, area 1, element 1, layer 3, skeleton 6/ DA228 (27) |  |  |  |  |
|  |  | Middle Ages (901-920 CE) | Kazakhstan / Almaly, Kurgan 1, Object 1, Issyk, Tian Shan / DA126 (27) |  |  |  |  |
| 9 | m.5628T>C, tRNA Alanine/L | Iron Age (400-200 BCE) | Moldova / Glinoe / SCY308 (59) | 0.19% | LP | Pathogenic (M2-P31) | CPEO / DEAF enhancer / gout (Heteroplasmy) |
| 10 | m.7543A>G tRNA Aspartic acid /H | Neolithic (8200-7700 BCE) | Iran / Tepe Abdul Hosein, Central Zagros / AH1 (60) | 0.09% | PP | Pathogenic (M2-P29) | MEPR (Heteroplasmy) |
| 11 | m.7554G>A, tRNA Aspartic acid/H | Neolithic (8205-7756 BCE) | Iran / Tepe Abdul Hosein, Central Zagros / AH2 (60) | 0.00% | PP | Pathogenic (M2-P40) | Myopathy, ataxia, nystagmus, migraines, lactic acidosis (Heteroplasmy) |
|  |  | Middle Ages (10-375 CE) | Poland / Kowalewko, Greater Poland / PCA0018 (55) |  |  |  |  |
| 12 | m.8296A>G tRNA Lysine/H | Bronze (2872-2583 BCE) | Russia / Grachevka, Sok River, Samara / I0371 (58) | 0.07% | PP | Pathogenic (M2-P2) | DMDF / MERRF / HCM / epilepsy (Heteroplasmy) |
| 13 | m.8328G>A tRNA Lysine/H | Middle Ages (80-260 CE) | Poland / Kowalewko, Greater Poland / PCA0054 (55) | 0.00% | LP | Pathogenic (M2-P39) | Mito Encephalopathy / EXIT with myopathy and ptosis (Heteroplasmy) |
| 14 | m.8342G>A<br>rs118192103/<br>tRNA-Lysine/H | Middle Ages<br>(10-375 CE) | Poland / Kowalewko, Greater<br>Poland / PCA0018 (55) | 0.00% | LP | Pathogenic<br>(M2-P53) | CPEO and Myoclonus<br>(Heteroplasmy) |
| 15 | m.12300G>A<br>tRNA<br>Leucine/H | Middle Ages<br>(80-260 CE) | Poland / Kowalewko, Greater<br>Poland / PCA0004 (55) | 0.00% | PP | Pathogenic<br>(M0-P36) | MERRF<br>(Heteroplasmy) |
| 16 | m.15915G>A<br>tRNA<br>Threonine/H | Middle Ages<br>(80-120 CE) | Poland / Kowalewko, Greater<br>Poland / PCA0032 (55) | 0.00% | PP | Pathogenic<br>(M2-P30) | Encephalomyopathy<br>(Heteroplasmy) |

201B Freq –GenBank frequencies; CPEO - Chronic progressive external ophthalmoplegia; MM - Mitochondrial myopathies; MERRF - Myoclonic epilepsy 210with ragged red fibers; MEPR - Myoclonic epilepsy and psychomotor regression; DMDF - Diabetes Mellitus + Deafness. HCM - Hypertrophic 211cardiomyopathy; MERRF – Myoclonic epilepsy with ragged red fibers; MEPR - Myoclonic epilepsy and psychomotor regression; MELAS - Mitochondrial 212Encephalomyopathy, lactic acidosis. and stroke-like episodes; MIDD - Maternally inherited diabetes and deafness; SNHL - Sensorineural hearing loss; NARP 213Neuropathy, ataxia and retinitis pigmentosa; Cfm Path – confirmed pathogenic; LP – likely pathogenic; PP – possibly pathogenic; M - tRNA model; P – 214position in tRNA model (HmtVar).

**Figure 1.**
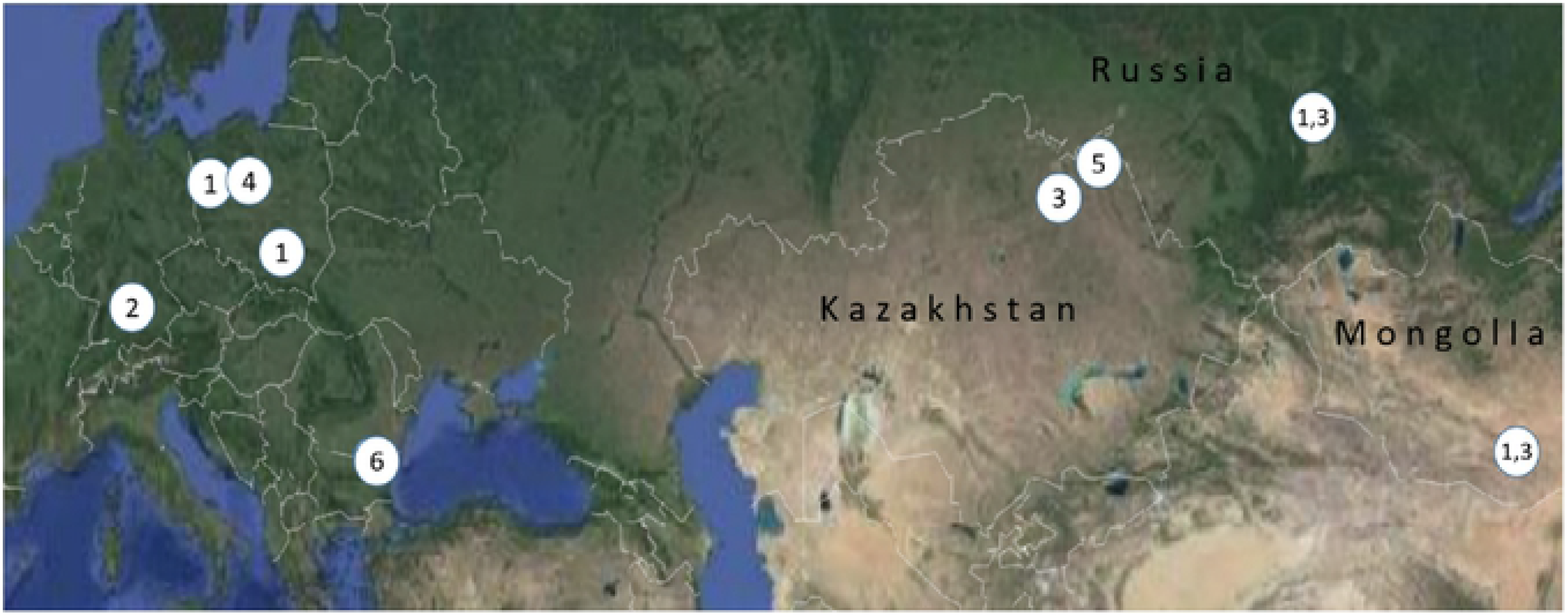
Geographic location of the Eurasian archaeological sites where mtDNA samples containing “confirmed pathogenic” variants had been found (cf. Table 1). Map source: Google Maps

Our literature survey for the m.5703G>A mutation finds that it has been reported to cause mitochondrial myopathy and ophtalmoplegia (MM) (28, 29). Recently, its phenotypic spectrum was broadened by a report of a patient with typical myoclonic epilepsy with ragged red fiber (MERRF) syndrome carrying a heteroplasmic m.5703G>A mutation (30, 31). In another recent study, however, it has been argued that investigations carried out to confirm pathogenicity of this mutation are insufficient (32).

#### m.3243A>G (rs199474657)

This variant was detected in a sample from a site in Germany from the Bronze Age (2029-1911 BCE) (Table 1, Figure 1). The mutation is in the *MT-TL1* gene, which encodes tRNA Leucine. MitoMap and HmtVar designate this mutation as pathogenic.

Population-based studies suggest the m.3243A>G mutation is the most common disease-causing mtDNA mutation, with a carrier rate of 1 in 400 people (33). The pathogenic mitochondrial DNA m.3243A>G mutation has been shown to be associated with a wide range of symptoms (34). Elevated heteroplasmy at this mtDNA site has been shown to lead to neurologic, sensory, movement, metabolic, and cardiopulmonary impairments (35). This mutation is associated with mitochondrial encephalopathy lactic acidosis and stroke-like episodes (MELAS) (36), maternally inherited deafness and diabetes (MIDD) (37) and chronic progressive external ophthalmoplegia (CPEO) (38). Other reported features include renal failure (39), isolated myopathy, cardiomyopathy, seizures, migraine, ataxia, cognitive impairment, bowel dysmotility and short stature (40). Low to moderate levels of mutant heteroplasmy in the m.3243G>A mutation are often associated with MIDD, whereas higher levels are variably associated with myopathy, high frequency sensorineural hearing loss, short stature, epilepsy, strokes and dementia (11, 41, 42).

#### m.5650G>A

This variants was found in 3 samples from the Iron Age, but they are from different Central Asian sites and time periods (ranging from 900 BCE to 134 BCE), so the individuals they were taken from are in all likelihood unrelated. This mutation is in the *MT-TA* gene, which encodes tRNA Alanine. MitoMap and HmtVar designate this mutation as pathogenic (Table 1, Figure 1).

McFarland et al. (2008) report a family where proximal myopathy has become increasingly severe with successive generations of the maternal lineage, and this pure myopathy is shown to be caused by the m.5650G>A mutation (43). Finnila et al. (2001) describe a patient with MERRF, who had a *de novo* m. 5650G>A mutation in the tRNA Alanine gene (44). The mutation was heteroplasmic, with the proportions of the mutant genome being up to 99% in muscle. They suggest that the mtDNA mutation is pathogenic as it was associated with a relevant clinical phenotype, it is absent in controls, and it alters a structurally important segment in the amino acid acceptor stem in the tRNA Alanine (Figure 2).

**Figure 2.**
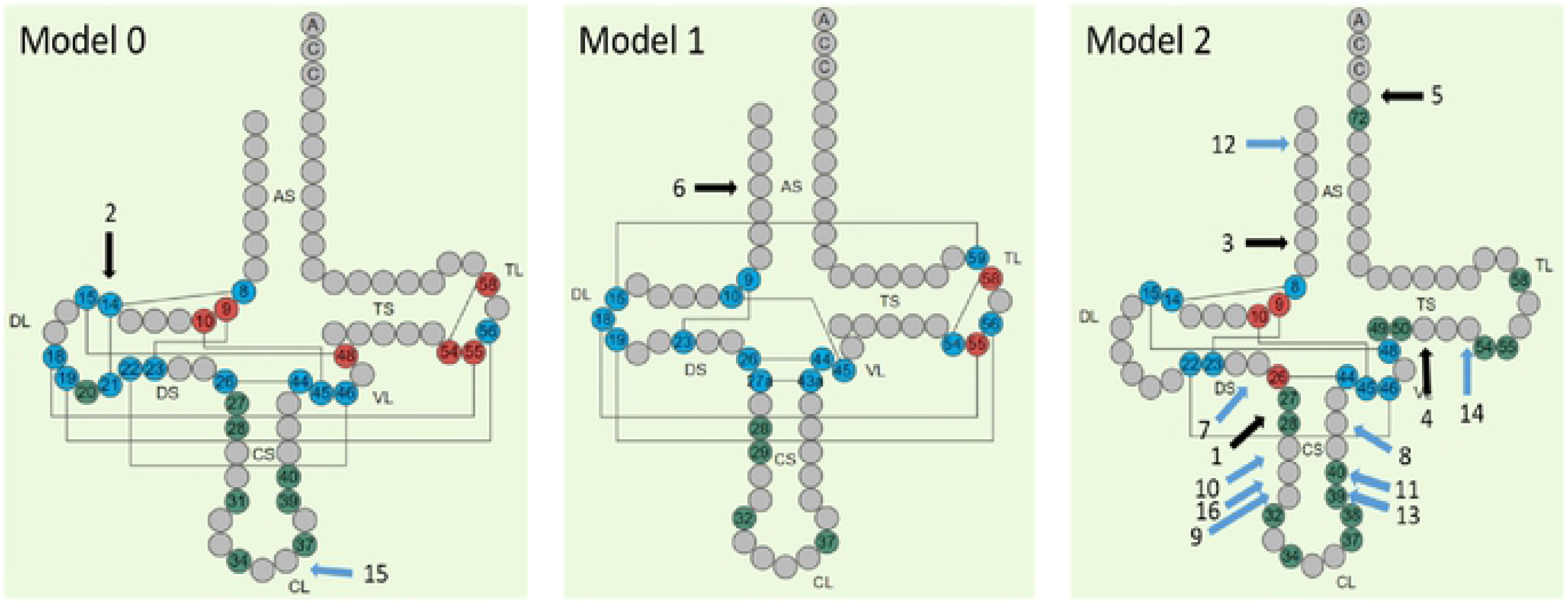
Location on tRNA nucleotide sequence of ancient Cfrm/LP/PP mutations (HmtVar). The positions of the tRNA nucleotide sequence affected by mutations are indicated with a black arrows for cfrm mutations and blue arrows for LP/PP mutations.

#### m.8340G>A

The variant was found in one sample from the Middle Ages (10-375 CE). This mutation is in the *MT-TK* gene, which encodes tRNA Lysine. MitoMap and HmtVar designate this mutation as pathogenic.

A number of studies have established association between the m.8340G>A variant and various clinical conditions. Jeppesen et al. (2014) report two patients with a *de novo* m.8340G>A variant associated with exercise intolerance, CPEO and myopathy (45). Gill et al (2017) report a patient carrying this mutation with cataracts, pigmented retinopathy, rod-cone dysfunction and sensory neural deafness without myopathy (46). Tamopolsky et al. (2019) report a case with a *de novo* m.8340G>A DNA mutation associated with mitochondrial myopathy, ptosis and ophthalmoparesis, corroborating the pathogenicity of the m.8340G>A mutation (47). The collective data is consistent with a categorization of “pathogenic” given that the variant is found in much higher frequency in patients (3 reports) *vs* a control population and with much higher heteroplasmy levels reported in ragged blue fibers and COX-negative *vs* healthy fibers (45, 46).

#### m.14674T>C

The variant was found in one sample from Poland from the Bronze Age (1592-1591 BCE). This mutation is in the *MT-TE* gene, which encodes tRNA Glutamic acid. MitoMap and HmtVar designate this mutation as “pathogenic*”* (Table 1, Figure 1).

A study by Houshmand et al (1994) describes two patients with Mitochondrial myopathy (MM) that are carriers of the m.14674T>C mutation. Even though it is absent in controls, the authors presume this mutation, as it does not change nucleotides that are conserved between species, is not likely to be pathogenic. This homoplasmic mutation has been identified in reversible infantile cytochrome c oxidase deficiency (or “Benign COX deficiency”) (48). Carriers of this mutation experience subacute onset of profound hypotonia, feeding difficulties and lactic acidosis within the first months of life. Although recovery may occur, mild myopathy persists into adulthood (49).

#### m.7510T>C (rs199474820)

This variants was found in one sample from Bulgaria from the Neolithic period, estimated to be from 5800-5400 BCE, and is thus the oldest pathogenic variant detected in this study. This mutation is located in the *MT-TS1* gene, which encodes tRNA Serine. MitoMap and HmtVar designates this mutation as pathogenic (Table 1, Figure 1).

A number of studies have established association between the m.7510T>C mutation and non-syndromic sensorineural hearing impairment (SNHL) (50–53). Review of the published cases suggests that there is interfamilial variability in the age of onset, accompanying symptoms, and haplogroup background (50). The results of Kytövuori *et al* (2017) suggest that in addition to sensorineural hearing impairment, the m.7510T>C mutation is associated with a spectrum of mitochondrial disease clinical features including migraine, epilepsy, cognitive impairment, ataxia, and tremor, and with evidence of mitochondrial myopathy (54).

### Likely/possibly pathogenic mutations

This group includes mutations for which there is discrepancy in their clinical effect designation between the two used platforms. Whereas HmtVar classifies them as pathogenic, MitoMap classifies them as likely or possibly pathogenic (Table 1).

## Discussion

This study establishes for the first time the presence of pathogenic mtDNA variants in 1443 ancient mtDNA Eurasian genomes from different periods, publicly available in the AmtDB database. Among the 3191 unique variants in the analyzed samples six are “confirmed pathogenic” with well-established clinical association in the contemporary patients.

The established “*confirmed pathogenic*” mutations are detected in samples estimated to be between 1600 and 7800 years old. The oldest of these, m.7510T>C, is detected in a Neolithic sample (5800-5400 ВСЕ). The m.5703G>A mutation was detected in a Neolithic sample that could be as old as 4800 years, but it is found again in samples from later periods, Iron Age (900-600 BCE) and Middle Age (10-375 CE). Other mutations with established long histories, detected in >4000 years old Bronze Age samples are m.3243A>G “MELAS”, the most common mtDNA mutation with pathogenic effect, present in contemporary populations with 0.02% frequency, and m.14674T>C, found in a 2600 old Bronze Age sample, and also found in in contemporary populations with 0.01% frequency (cf. Table 1). The m.5650G>A mutation is detected in an Iron Age sample that could be as old as 2900 years. The low population frequency of these variants in contemporary populations is further corroboration of their putative pathogenic significance.

Two mutations with likely/possibly pathogenic effect, m.7543A>G and m.7554G>A, in the *MT-TD* gene (tRNA Aspartic acid), detected in Neolithic period samples from the Neolithic period, from the same archeological site and time span, are the oldest established mutations with putatively pathogenic effect (10200-10000 years old). Also, two Bronze Age mutations, m.8296A>G and m.4440G>A, could be as old as 4800 years. It is noteworthy that one of these, m.4440G>A, is detected in as many as 12 ancient samples from different time periods and locations. Its pathogenic effect is corroborated by that it is the most common clinically significant mutation detected in ancient mtDNA samples in this study, yet it has been described as a novel mutation causing mitochondrial myopathies in contemporary patients (61). The remaining mutations in this category are detected in samples from later periods (2200-2000 years old).

It is noticeable that all the established ancient mutations putatively associated with diseases are located in tRNA genes, and none is in genes encoding the 13 essential polypeptides of the OXPHOS system, even though tRNAs comprise only about 10% of the total coding capacity of the mitochondrial genome (62). Epidemiological studies have highlighted that point mutations in the mt-tRNA genes are among the most common defects observed (63, 64). Mitochondrial tRNA mutations have been shown to be the most prevalent genetic defect by a survey of an adult population with mtDNA disease, accounting for more than 50% of all genetically diagnosed cases (65). More than 150 different point mutations have been described in mt-tRNA genes including novel disease-causing mutations and associated pathogenic mechanisms continue to be identified (66), yet mtRNA mutations` role in interfering with the translation mechanism remains unclear.

Ancient mutations putatively associated with mitochondrial diseases are in different tRNA genes and affect nucleotides in different functional parts of the encoded tRNA molecule (Table 2).

**Table 2.**
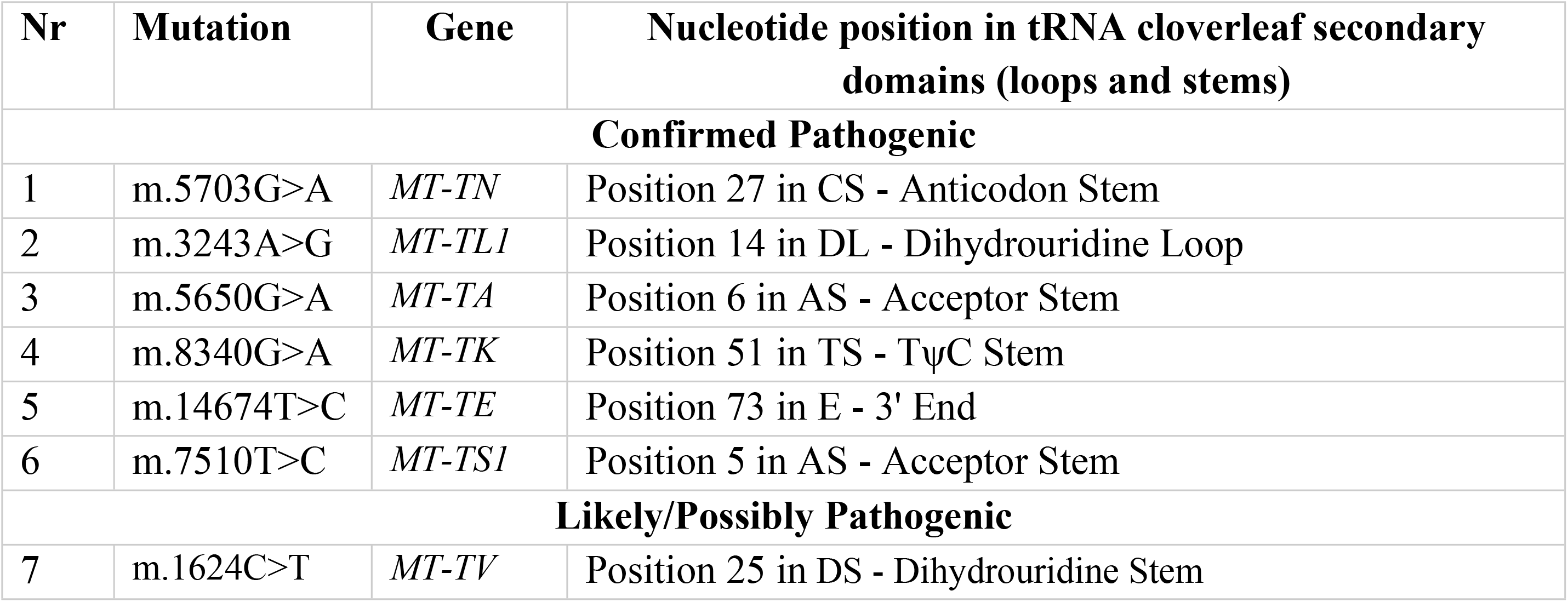
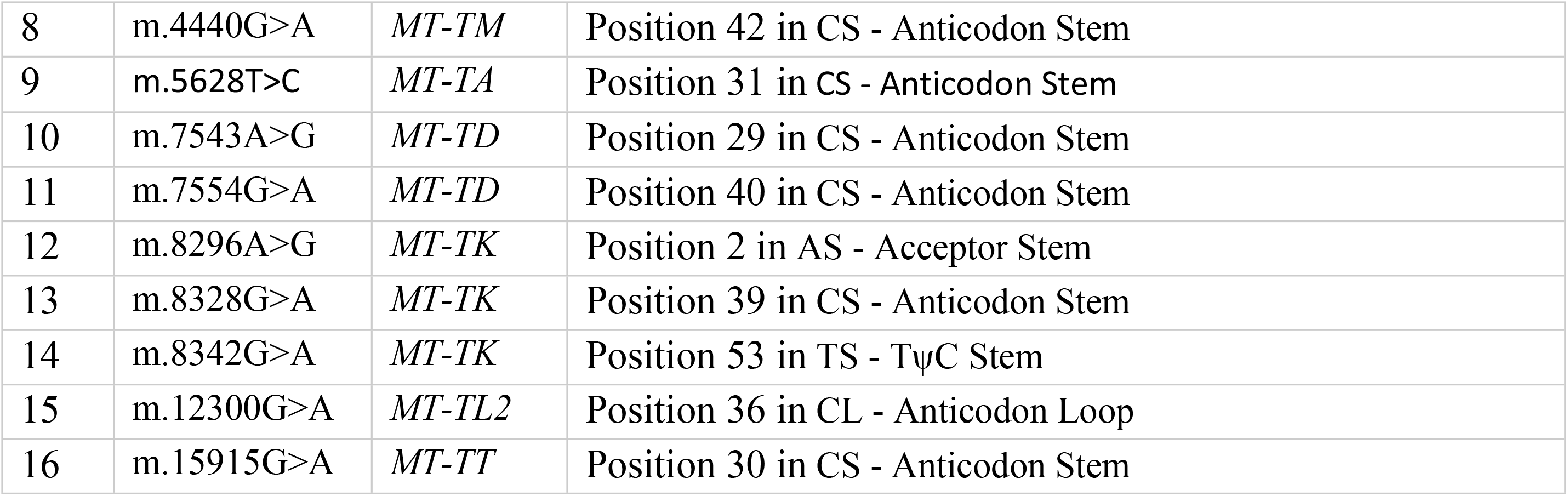
Nucleotide position in tRNA cloverleaf secondary domains of ancient mutations in tRNAs

The gene *MT-TK* coding tRNA Lysine is most commonly affected by mutations putatively associated with mitochondrial diseases in ancient mtDNA samples, i.e. one confirmed pathogenic (m.8340G>A) and three likely/possibly pathogenic (m.8296A>G, m.8328G>A and m.8342G>A). It is conceivable that these mutations have different clinical significance related to their impact on the stability of the tRNA protein.

A recent study on another mtDNA mutation in the same gene - m.8344 A>G, known to be associated with MERRF, has demonstrated that tRNA modifications have distinct effects on the stability and synthesis of mitochondrial proteins (45). Such regulating mechanisms conceivably contribute to human disease, and new RNA sequencing approaches to mitochondria should provide insights. Two ancient mutations were established in the *MT-TD* coding Aspartic acid. Aspartic acid is neurotransmitter and recent studies show that it may be involved in the pathogenesis of a stroke-like episode (46). The remaining putatively pathogenic mutations established in ancient mtDNA are located in different tRNA genes.

It is noteworthy that the established ancient mutations are located in different cloverleaf models of tRNAs. (Figure 2). The tRNA model indicates one of the four possible groups human mt-tRNAs are classified based on their structural diversity and tertiary interactions (67). Model 0 model represents the quasi-canonical cloverleaf structure, with standard D-loop/T-loop interaction; Model 1-a single tRNA with an atypical anticodon stem; Model 2 – the most common among mt-RNAs, is characterized by loss of D-loop/T-loop interaction and and Model 3 - lack of D-stem. Thirteen out of the sixteen mutations putatively associated with mitochondrial disease presented in this study are in Model 2.

Seven out of the 16 mutations considered here are located in CS-Anticodon Stem, four in AS-Acceptor Stem, two in TS-TΨC Stem, and single mutations are located in DL-Dihydrouridine Loop, CL-Anticodon Loop and DS-Dihydrouridine Stem (Fig. 2).

Confirmed pathogenic mutation m.5703G>A is located on position 27 in Model 2 tRNA which is involved in post-transcriptional modifications and is adjacent to position 26, which participates in tertiary folding and is also subject to posttranscriptional modifications. Confirmed pathogenic mutation m.3243A>G in Model 0 t RNA affects position 14, which is involved in tertiary folding with interactions represented by lines on Fig. 2. Mutation m.14674T>C in Model 2 tRNA is in position 73 in 3’ End of the acceptor stem, which participates in 3’ end-editing in the final stages of tRNA formation. The role of the remaining three confirmed pathogenic mutations is more difficult to be determined. Mutation m.5650G>A is in position 6 of the acceptor stem of a Model 2 tRNA. Pathogenic mutation m.8340G>A in position 51 of a Model 2 tRNA is located next to position 50 nucleotide involved in post-transcriptional modifications.

Three mutations with putatively pathogenic effect are in positions that affect the structure and the function of the tRNA, m.12300G>A in position 36 of the CL-Anticodon Loop, m.8328G>A in position 39 and m.7554G>A in position 40 that all have impact on post-transcriptional modifications. Mutations in tRNA genes could lead to disturbances of three dimensional structure, the absence of post-transcriptional modifications of the tRNA, rise in the number of errors and thus tRNA destabilization.

These changes could have an effect on the codon decoding speed, leading to accumulation of cell damaging proteins and cutback of mitochondrial protein synthesis, including reduction of OXPHOS proteins and insufficiency of the OXPHOS constituent complexes I, III, and IV. Despite the clinical significance, the molecular mechanisms leading to such disturbances remain poorly understood.

As is often happens in ancient DNA analyses, because the amount of endogenous template DNA is typically very low, the surviving molecules are typically short and affected by post-mortem cytosine deamination damage which appears as C>T and G>A variants in sequence data (68). It is noteworthy to mention that 12 out of the 19 (63.2%) pathogenic or putatively pathogenic mutations that we detect in the analyzed ancient mtDNA samples are G>A or C>T substitutions. Review of the publications that present the analyzed ancient mtDNA genomes substantiates that the authors have employed adequate analyses to mitigate the effect of post mortem damage (PMD), strengthening our confidence that these are real variants and not the result of PMD (27, 69, 70).

Still, pathogenic mutations in mitochondrial DNA, often show highly variable phenotypes for any given point mutation and severity of the clinical and biochemical phenotype has been roughly proportionate to the percent mutant heteroplasmy (10, 41). Identifying heteroplasmic variants and establishing the level of heteroplasmy in ancient samples is not a trivial task. Heteroplasmic variants however constitute the bulk of disease-associated mtDNA variants in contemporary humans, and most of the detected pathogenic or putatively pathogenic variants in our study have pathogenic effect in heteroplasmic state. Nevertheless due to insufficient phenotypic data about the human remains, there is no way of exactly knowing if disease-associated mutations, or those predicted to have a strong functional effect, were indeed pathogenic in ancient populations.

## Conclusion

The established mtDNA pathogenic mutations in the analyzed ancient samples are putatively associated with a wide range of mitochondrial diseases found in contemporary populations. Studying putative pathogenic mutations from ancient mtDNA informs on the mitochondrial disease spectrum in ancient times, and comparing their frequencies among populations separated by significant time periods sheds light on the history of the disease. Our findings suggest that disease associated genes are often genes with long history rather than newly evolved genes (71), warranting further research attention. The dynamics of the prevalence of putative pathogenic variants in paleogenetic and contemporary genetic data can be used to predict the future course of human microevolution.

## Contributions

D.T. – conceived the study, analyzed results, discussed data and wrote the paper; D.S., S.K.-Y.,

D.N. – analyzed results, discussed data. All authors reviewed the manuscript.

## Corresponding author

Correspondence to

## Competing interests

The authors declare no competing interests.

